# Quercetin reduces APP expression, oxidative stress and mitochondrial dysfunction in the N2a/APPswe cells via ERK1/2 and AKT pathways

**DOI:** 10.1101/2022.09.18.508406

**Authors:** Zhi Tang, Min Guo, Yaqian Peng, Ting Zhang, Yan Xiao, Ruiqing Ni, Xiaolan Qi

**Affiliations:** Key Laboratory of Endemic and Ethnic Diseases, Ministry of Education and Key Laboratory of Medical Molecular Biology of Guizhou Province, Guizhou Medical University, Guiyang, China; Collaborative Innovation Center for Prevention and Control of Endemic and Ethnic Regional Diseases Coconstructed by the Province and Ministry, Guiyang, China; Institute for Regenerative Medicine, University of Zurich, Zurich, Switzerland; Institute for Biomedical Engineering, ETH Zurich and University of Zurich, Zurich, Switzerland

**Author notes:** These authors equally contributed to this manuscript (Zhi Tang, Min Guo). To whom correspondence should be addressed: Ruiqing Ni,; Xiaolan Qi,.

**Keywords:** AKT pathway, APP, ERK1/2 pathway, mitochondrial dysfunction, oxidative stress, quercetin

## Abstract

Abnormal amyloid-β (Aβ) abnormal accumulation and oxidative stress play important roles in Alzheimer’s disease (AD). Quercetin has been reported to possess antioxidant and anti-inflammatory properties, and thus of therapeutic interests for neurodegenerative disorders. In the present study, we aimed to characterize the mechanisms by which quercetin exerts neuroprotective effects in murine neuroblastoma N2a cells stably expressing human Swedishh mutant amyloid precursor protein (APP). Quercetin treatment exhibited low cytoxicity, attenuated APP expression and APP-induced oxidative neurotoxicity in N2a/APP cells. We found that quercetin effected via the down-regulation of phospho-extracellular signal□regulated protein kinase (p-ERK1/2) pathway and up-regulation of phospho-protein kinase B (p-AKT) pathway in N2a/APP cells. In addition, quercetin ameliorated the elevated levels of reactive oxygen species using DCFH-DA flow-cytometry in N2a/APP cells, lipid peroxidation using (4-HNE), and DNA oxidation (8-OHdG assays). Quercetin ameliorated the loss of mitochondrial membrane potential using JC-1 fluorescence assay in N2a/APP cells in a dose-dependent mannor. In conclusion, we domenstrated the neuroprotective effects of quercetin against the APP expression induced oxidative neurotoxicity, impairment of mitochondrial function and oxidative stress through inactivation of the ERK1/2 signaling pathway and activation of AKT signaling pathways.

## 1. Introduction

Alzheimer’s disease (AD) is the most common neurodegenerative disorder occurring mainly in elderly individuals, clinically characterized by progressive memory deficits and cognitive dysfunction (Scheltens et al., 2021). The neuropathological hallmarks of AD include senile plaques, neurofibrillary tangles and neuronal loss (Braak and Braak, 1991). The main component of senile plaques is the β-amyloid (Aβ) peptide, which is derived from amyloid precursor protein (APP) via proteolysis by β-secretase and γ-secretase (Selkoe and Hardy, 2016). Aβ is neurotoxic and its abnormal accumulation leads to oxidative stress, glial cell activation and neuronal loss (Lempriere, 2021; Mucke and Selkoe, 2012). Therefore, blocking Aβ production and aggregation has been recognized as one of the most crucial strategies for preventing and treating AD (Panza et al., 2019). Increased oxidative stress has been proposed as one of the possible common aetiologies in AD (Butterfield and Halliwell, 2019; Su et al., 2008). Chronic oxidative stress could induce DNA repair system impairment, cellular damage, and mitochondrial dysfunction (Niedzielska et al., 2016). Furthermore, oxidative stress could in turn provoke Aβ production and aggregation, which could lead to a vicious pathogenic cycle in the pathogenesis of AD (Fyfe, 2022; Ionescu-Tucker and Cotman, 2021).

Quercetin (3,3,4,5,7-pentahydroxyflavone) is one of the major flavonoids that is present in a variety of vegetables and fruits (Singh et al., 2021). Quercetin exhibits a wide range of beneficial activities, such as antioxidant, anti-inflammatory, anti-proliferative and anti-apoptotic effects on various cells and animal models (Ebrahimpour et al., 2020; Farr et al., 2017; Islam et al., 2021; Novais et al., 2021; Suganthy et al., 2016). In addition, quercetin exhibit antiamyloidogenic, Aβ fibril-disaggregating (Jiménez-Aliaga et al., 2011), as well as anti-tau aggregation effects *in vivo* (Musi et al., 2018). Several studies have demonstrated the effect of quercetin in animal models of AD (Sabogal-Guaqueta et al., 2015; Zhang et al., 2019; Zhang et al., 2016). In addition, quercetin treatment has been previous reported in patients with early-stage AD (Nakagawa et al., 2016). Dasatinib plus quercetin traement is currently in phase 1/2 clinical trial in adults at risk for AD (NCT05422885) after the proof-of-concept SToMP-AD study (NCT04063124) (Gonzales et al., 2022).

The extracellular signal-regulated kinase 1/2 (ERK1/2), and the protein kinase B (AKT) pathways play pivotal roles in oxidative inbalance in AD (Cao et al., 2019; Rai et al., 2019), has been shown involved in mediating Aβ production via APP processing (Amanzadeh et al., 2019; Hefter et al., 2020; Rai et al., 2019). Aberrant accumulation of activated ERK1/2 and AKT in neurons has been reported in AD brains (Ferrer et al., 2001; Pei et al., 2002; Pei et al., 2003; Rai et al., 2019). In this study, we hypothesize that quercetin attenuates APP expression and oxidative stress, thereby providing a defensive mechanism against APP-induced oxidative neurotoxicity via ERK1/2 and AKT pathways.

## 2. Material and Methods

### 2.1. Materials, reagents and antibodies

Details of chemicals, antiboides and kits used in the study were listed in supplementary Table

### 2.2. Cell culture, treatment and extracts

N2a cells stably transfected with human APP Swedish mutant (N2a/APP) were gifts from Professor Rong Liu (Tongji Medical School, Wuhan, China). The cells were cultured in an equivalent volume of Dulbecco’s modified Eagle’s medium (DMEM, Gibco, USA) with 10% fetal bovine serum (Gibco, USA) and 1% penicillin□streptomycin solution (HyClone, USA) in 5% CO_2_ at 37 °C. The cells were grown to 80% confluence in 25 cm^2^ culture flasks and passaged into 6-well culture plates. After treatment with quercetin at concentrations of 10, 20 and 50 μM for 24 h, the cells were washed three times with ice-cold phosphate-buffered saline (PBS, pH 7.4, Biological Industries, Israel), harvested, and suspended in 100 μL RIPA cell lysis buffer (Solarbio, China) containing protease inhibitor cocktail (diluted 1:200, Sigma, USA). The cell lysates were sonicated on ice, and the protein concentration was determined with a Bradford kit (Bio-Rad, USA).

### 2.3. Cell viability assay

Cell viability was measured by a cell counting kit-8 (CCK-8) (MedChemExpress LLC, China) according to the manufacturer’s instructions. A total of 5×10^3^ cells were seeded into 96-well plates (Chen et al., 2022; Tang et al., 2021). After being treated, 10 μl of CCK-8 reagent and 100 μl of medium were added to each well, and incubated for 1 h at 37 °C. Next, the absorption (450 nm) was measured using a microplate reader (Bio-Rad, Hercules, USA).

### 2.4. Western blotting

Western blotting was performed as described previously (Ding et al., 2021; Lai et al., 2022). In brief, boiled samples were separated on 4%-12% (w/v) sodium dodecyl sulfate (SDS)–polyacrylamide gels (Absin, China), and the separated proteins were then transferred to polyvinylidene difluoride membranes (Millipore, USA). These membranes were blocked in 5% (w/v) nonfat milk (Coolaber, China) for 2 h and incubated with primary antibodies overnight at 4 °C. Subsequently, the membrane was washed three times with TBST buffere consisting of Tris-buffered saline (TBS, Servicebio, China) supplemented with 0.1% (v/v) Tween-20 (Solarbio, China), and the bound IgGs were recognized by anti-mouse or anti-rabbit goat anti-Mouse IgG (H+L), horseradish peroxidase (HRP) or goat anti-Rabbit IgG (H+L), HRP secondary antibodies (Thermo Fisher Scientific, USA). The complexes were visualized with a chemiluminescence imaging system (SYNGENE, GeneGnome XRQ NPC, UK). The band density was analysed by ImageJ software (NIH, USA).

### 2.5. Reactive oxygen species (ROS) detection by flow-cytometry

To detect intracellular ROS, the fluorescent probe 2,7-dichlorofluorescein diacetate (DCFH-DA, Beyotime, China) was used. Na/APP cells in 6-well plates at a density of 2 ×10^5^/well were treated without or with quercetin at the concentration of 10 μM, 20 μM, 50 μM for 24 h. Te cells were maintained in DMEM containing 10 μM DCFH-DA at 37 °C for 20 min. The staining solution containing DCFH-DA was removed afterwards. The cells were gently washed twice with DMEM. Then, the cell suspension was loaded into a flow-specific tube and detected by flow cytometry at excitation and emission wavelengths of 485 and 538 nm, respectively. The fluorescence was measured by BD FACSVerse™ flow-cytometry (BD Biosciences, USA). At least 10,000 events were acquired for further quantification. All data were analysed by using FlowJo software (BD Bioscience, USA).

### 2.6. Mitochondrial membrane potential detection

The mitochondrial membrane potential assay was performed to quantify the level of the polarization/depolarization of the mitochondrial membrane by using the JC-1 dye (Beyotime, China) after treatment with quercetin. N2a/APP cells were treated with quercetin (0, 10 μM, 20 μM, 50 μM) at a density of 2×10^5^ cells/well in a 6-well plate. After 24 h, the cells were collected and stained using JC-1 dye. Finally, the cells were analysed on a BD FACSVerseTM flow cytometer, and the results were analysed by using FlowJo software.

### 2.7. Lipid peroxidation and DNA damage marker measurement

The levels of 4-hydroxynonenal (4-HNE) produced by lipid peroxidation and 8-hydroxy-2’-deoxyguanosine (8-OHdG) as DNA damage marker after treatment with quercetin were assessed, N2a/APP cells were treated with quercetin (0, 10 μM, 20 μM, 50 μM) and plated onto polylysine-coated coverslips were fixed with 4% paraformaldehyde in PBS pH 7.4 for 15 min at room temperature. Cells were permeabilized by treatment with TBS containing 0.1% Triton X-100 (Solarbio, China) for 5 min and then incubated with primary antibodies against 4-HNE (1:150) and 8-OHdG (1:150) overnight at 4 °C. After washing with PBS pH 7.4, bound antibody was detected by incubation for 1 h with secondary antibodies conjugated with Alexa Fluor 488 (1:200, Thermo Fisher Scientific, USA) or Alexa Fluor 546 (1:200, Thermo Fisher Scientific, USA). Sections were mounted with vector medium containing 4’,6-diamidino-2-phenylindole (DAPI) Fluoromount-G (Vector laboratories, USA). All images were examined on an Olympus confocal microscope (Olympus, Japan) with a 40 × objective and a constant exposure time for each marker in all analysed sections. The experiments were repeated three times. A total of 100 cells from each group were analysed.

### 2.8. Statistical analysis

All analyses were performed with SPSS software, version 25.0 (IBM, USA) and GraphPad Prism version 9.0 (GraphPad, USA). The data are presented as the mean ± standard error (SEM) (n = 3 or 6). A total of 100 cells from each group were analysed. Statistical significance was determined by one-way ANOVA, followed by Dunnett’s multiple comparisons test with a 95% confidence interval. A p value ≤ 0.05 was considered statistically significant.

## 3. Results

### 3.1. Quercetin has low cytotoxicity for N2a/APP cells and reduces APP expression in N2a/APP cells

We first studied the cytotoxicity of quercetin on N2a/APP cells. No loss of cell viability was observed in N2a/APP cells treated for 24 h with 50 μM quercetin compared with controls treated without quercetin (0 μM). Treatment using 100 μM quercetin decreased the viability of N2a/APP cells (p = 0.0214, treated vs. control group) (**Fig. 1A**). Therefore, treatment for 24 h with a concentration range of 0~50 μM quercetin was used in the following investigations. Next we examined whether quercetin treatment could inhibit APP expression in N2a/APP cells. We found that the expression levels of APP were reduced after 24 h treatment with 50 μM quercetin in N2a/APP cells (p = 0.0241, treated vs. control group) based on the immunoblotting results (**Figs. 1B, C**).

**Figure 1.**
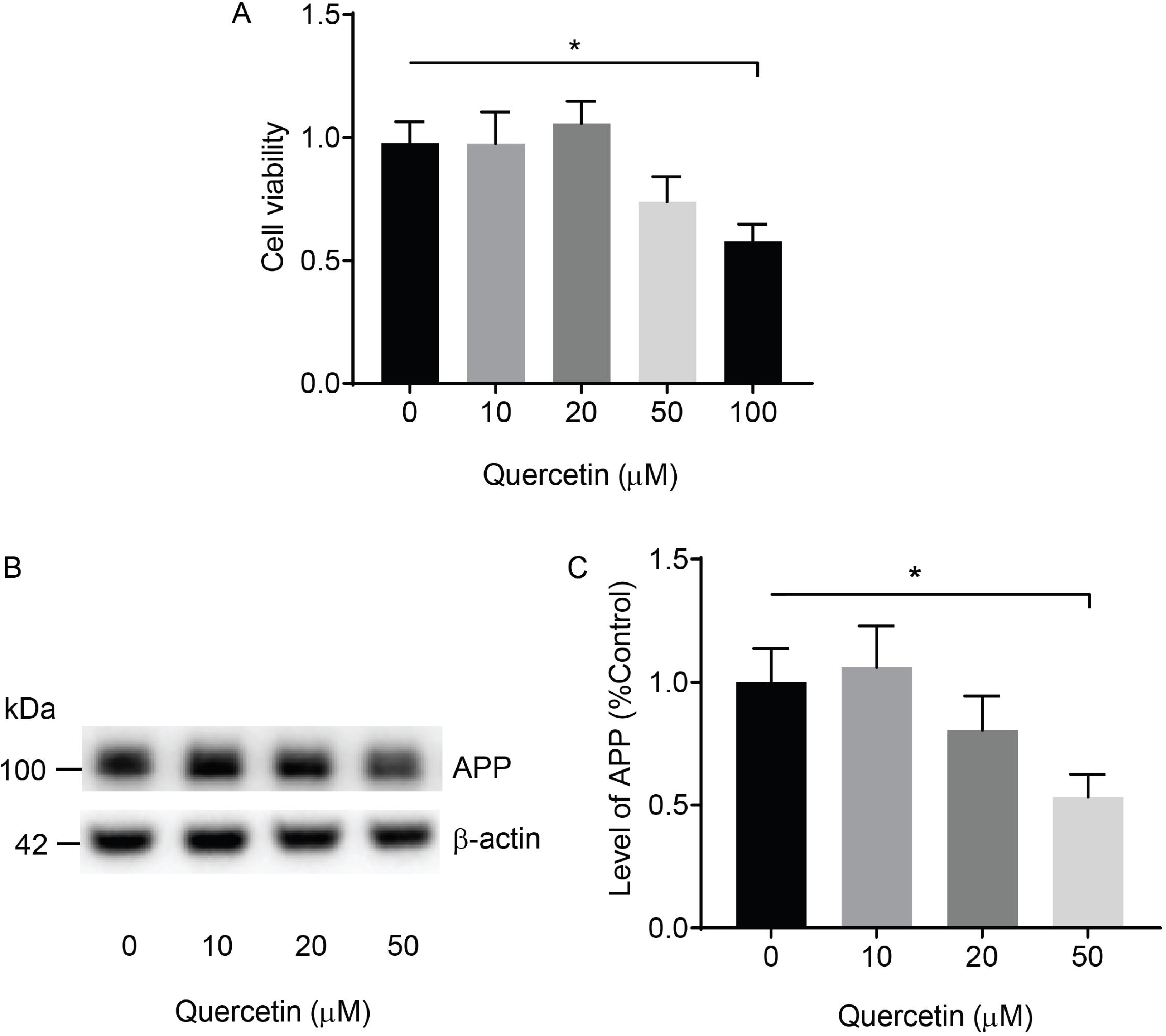
Quercetin exhibits low cytotoxicity and attenuates APP expression in N2a/APP cells. A, Cell viability was assessed using the CCK-8 assay. N2a/APP cells were treated without and with quercetin at the concentration of 10 μM, 20 μM, 50 μM for 24 h. Data are presented as the mean ± SEM (n = 6). *p < 0.05, one-way ANOVA, Dunnett’s multiple comparison. B, Representative immunoblots of APP proteins in N2a/APP cells in the absence or presence of quercetin at the concentration of 10 μM, 20 μM, 50 μM; C, Quantitation of the immunoblots in B (n = 3), *p < 0.05, one-way ANOVA, Dunnett’s multiple comparison.

### 3.2. Downregulation of ERK1/2 and upregulation of AKT pathways are involved in the protective effect of quercetin in N2a/APP cells

We hypothesized that the ERK1/2 and AKT signaling pathways are involved in effect of quercetin on Aβ production and aggregation. Then we assessed the protein levels of ERK1/2 and AKT following 24 h treatment with quercetin at concentrations of 10 μM, 20 μM and 50 μM in N2a/APP cells. Our data showed that the level of phosphorylated-ERK1/2 (Thr202/Tyr204, p-ERK1/2) was markedly decreased after 50 μM quercetin treatments (by 65 %, p = 0.0004, treated vs. control group), but not at lower concentration (**Figs. 2A, C**). In contrast, the levels of total-ERK1/2 (t-ERK1/2) remained unchanged compared to the control group after 50 μM quercetin treatments (**Figs. 2B, D**). Furthermore, the phosphorylation levels of AKT (p-ATK) at Ser473 were increased in the 50 μM quercetin-treated N2a/APP cells (by 20 %, p = 0.0429, treated vs. control group), but not at lower concentration (**Figs. 2E, G**). In contrast the levels of total-AKT (t-AKT) were not significantly different from those in control N2a/APP group (**Figs. 2F, H**). Thus, the protective effects of quercetin were associated with inactivation of ERK1/2 and activation of the AKT signaling pathways as defensive responses to oxidative stress.

**Figure 2.**
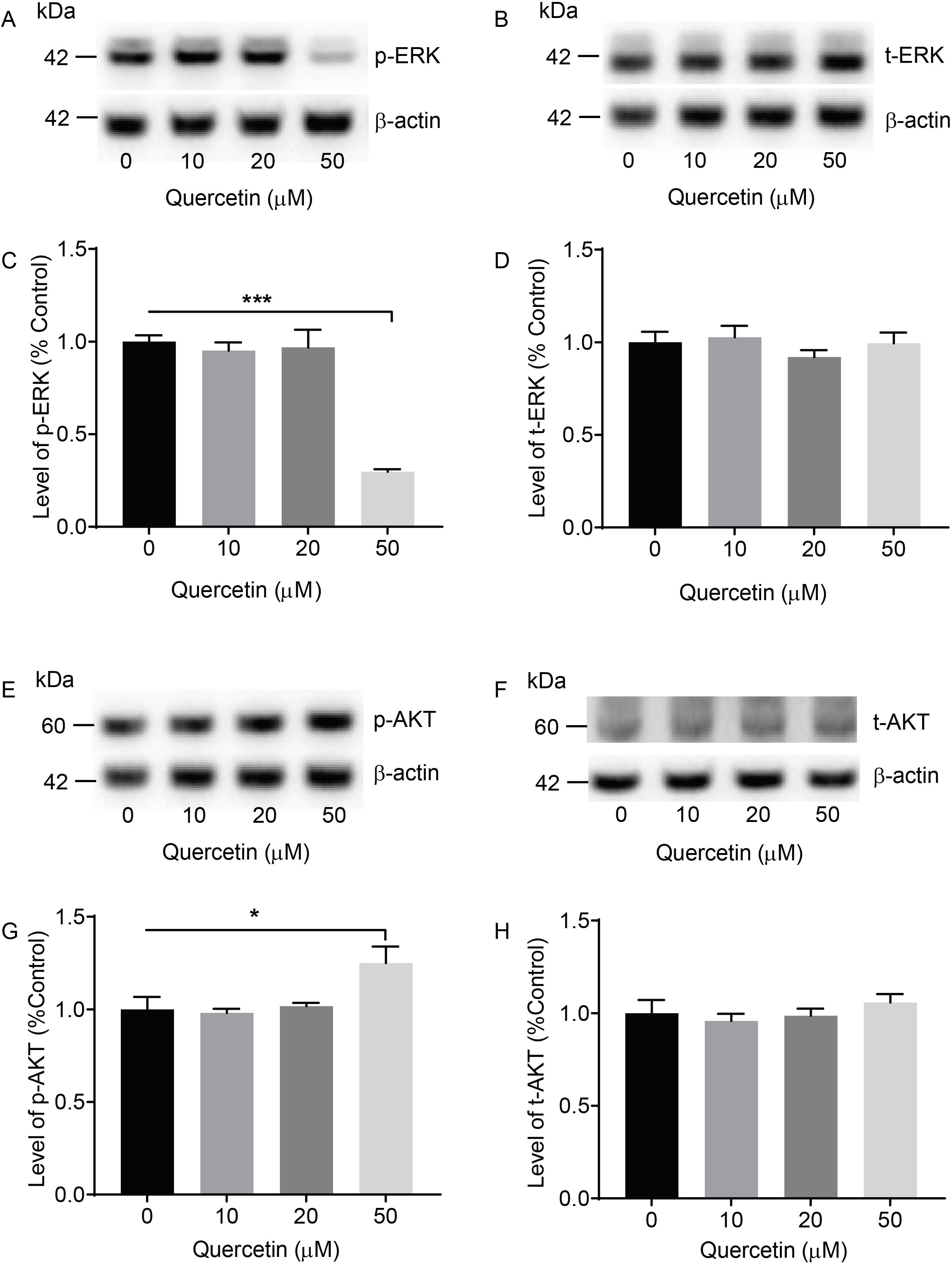
Quercetin attenuates the activation of the ERK1/2 pathways and enhances the activation of the AKT pathways in N2a/APP cells. A-B, Representative immunoblots of phosphorylated total-extracellular signal-regulated kinase (p-ERK1/2) and total ERK (t-ERK1/2) in N2a/APP cells without or with quercetin treatment at the concentration of 10 μM, 20 μM, 50 μM; ***p < 0.001, one-way ANOVA, Dunnett’s multiple comparison. C-D, Quantitation of the immunoblots in A, B (n = 3). E-F, Representative immunoblots of p-AKT and total AKT in N2a/APP cells with or without quercetin treatment at the concentration of 10 μM, 20 μM, 50 μM; G-H, Quantitation of the immunoblots in E, F (n = 3), *p < 0.05, one-way ANOVA, Dunnett’s multiple comparison.

### 3.3. Quercetin reestablished the loss of mitochondrial membrane potential ΔΨm) in N2a/APP cells

Maintaining Ψm is essential to ensure the scavenging efficiency of ROS and to confer cytoprotection towards apoptotic events caused by excessive ROS. Next, we investigated whether quercetin can ameliorate the mitochondrial dysfunction in N2a/APP cells. We measured the ΔΨm in N2a/APP cells by using JC-1 staining. JC-1 aggregates in the mitochondrial matrix and exhibits red fluorescence in healthy cells. When Ψm is diminished, JC-1 is converted to the monomer state, showing green fluorescence. The effects of quercetin on the mitochondrial membrane potential of N2a/APP cells were then examined using the ratio of green/red fluorescence. In the control N2a/APP cells group, a reduction in the Ψm after incubation was observed (green/red fluorescence ratio = 1.5)(**Figs. 3A, B**). Treatment with 50 μM quercetin markedly increased the Ψm, as indicated by a decrease in the ratio of green/red fluorescence intensity in N2a/APP cells by 65% (p < 0.0001, treated vs. control group, **Figs. 3A, B**). These findings suggested that quercetin could exert beneficial effects on mitochondrial function in N2a/APP cells.

**Figure 3.**
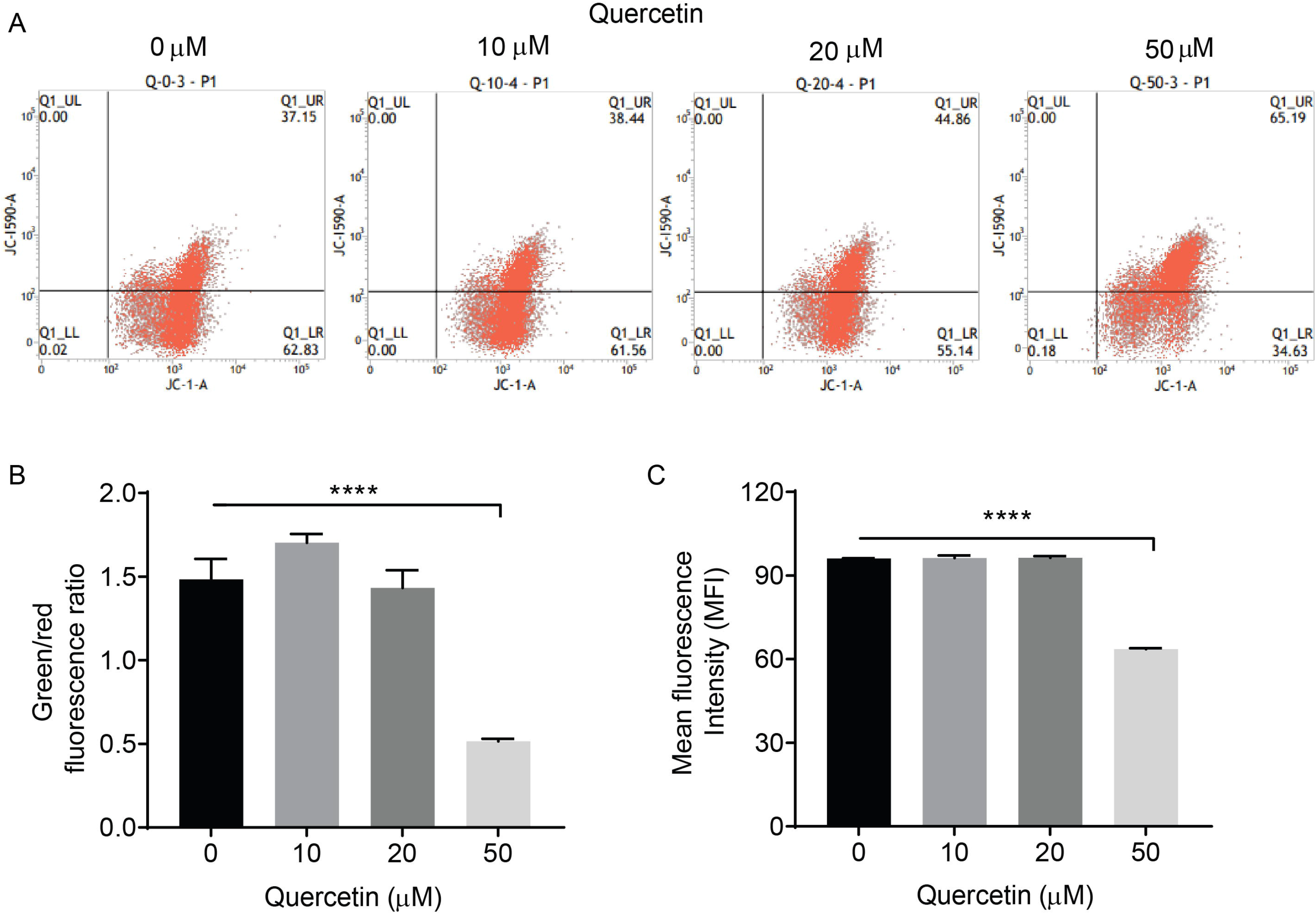
Quercetin reverses ΔΨm depolarization and the generation of ROS in N2a/APP cells. A, Analysis of ΔΨm with JC-1 staining by flow cytometry on N2a/APP cells without or with quercetin treatment at the concentration of 10 μM, 20 μM, 50 μM; B, The ratio of green fluorescence to red fluorescence (n = 3). ****p < 0.0001. C, Mean fluorescence intensity analysis of ROS with DCFH-DA staining by flow cytometry in N2a/APP cells without or with quercetin treatment at the concentration of 10 μM, 20 μM, 50 μM (n = 3). ****p < 0.0001, one-way ANOVA, Dunnett’s multiple comparison.

### 3.4. Quercetin prevents the generation of ROS in N2a/APP cells, thereby attenuating oxidative stress

Next we assessed the effect of quercetin at the concentration of 10 μM, 20 μM, 50 μM on ROS in N2a/APP cells by using the fluorescent DCFH-DA staining. We observed that the level of ROS (indicated by mean fluorescence intensity) in N2a/APP cells increased markedly after incubation period. Treatment with 50 μM quercetin counteracted this increase of the level of ROS, indicated by mean fluorescence intensity by 40 % (p < 0.0001, treated vs. control group, **Fig. 3C**), but not at lower concentration.

Next we assess the effect of quercetin on reducing the lipid peroxidation and DNA damage in N2a/APP cells by using 4-HNE and 8-OHdG assays in N2a/APP cells, respectively. The red fluorescence of 4-HNE produced by lipid peroxidation and the green fluorescence of stained 8-OHdG resulting from oxidation of DNA were elevated in N2a/APP cells after incubation in control group. These increases in 4-HNE and 8-OHdG fluroesence were both suppressed by quercetin treatment in a dose-dependent manner (**Figs. 4A-D**). Treatment with quercetin significantly reduced the fluorescence intensity of 4-HNE in N2a/APP cells at 20 μM (p = 0.0001, treated vs. control group) and 50 μM (p < 0.0001, vs. treated vs. control group) by xxx % and by xxx %, respectively (**Figs. 4A-B**). In addition, treatment with quercetin significantly reduced the fluorescence intensity of 8-OHdG in N2a/APP cells at 20 μM (p = 0.0001, treated vs. control group) and 50 μM (p < 0.0001, treated vs. control group) by xxx % and by xxx %, respectively (**Figs. 4C-D**). In summary, these observations indicated that quercetin ameliorated oxidative stress in N2a/APP cells.

**Figure 4.**
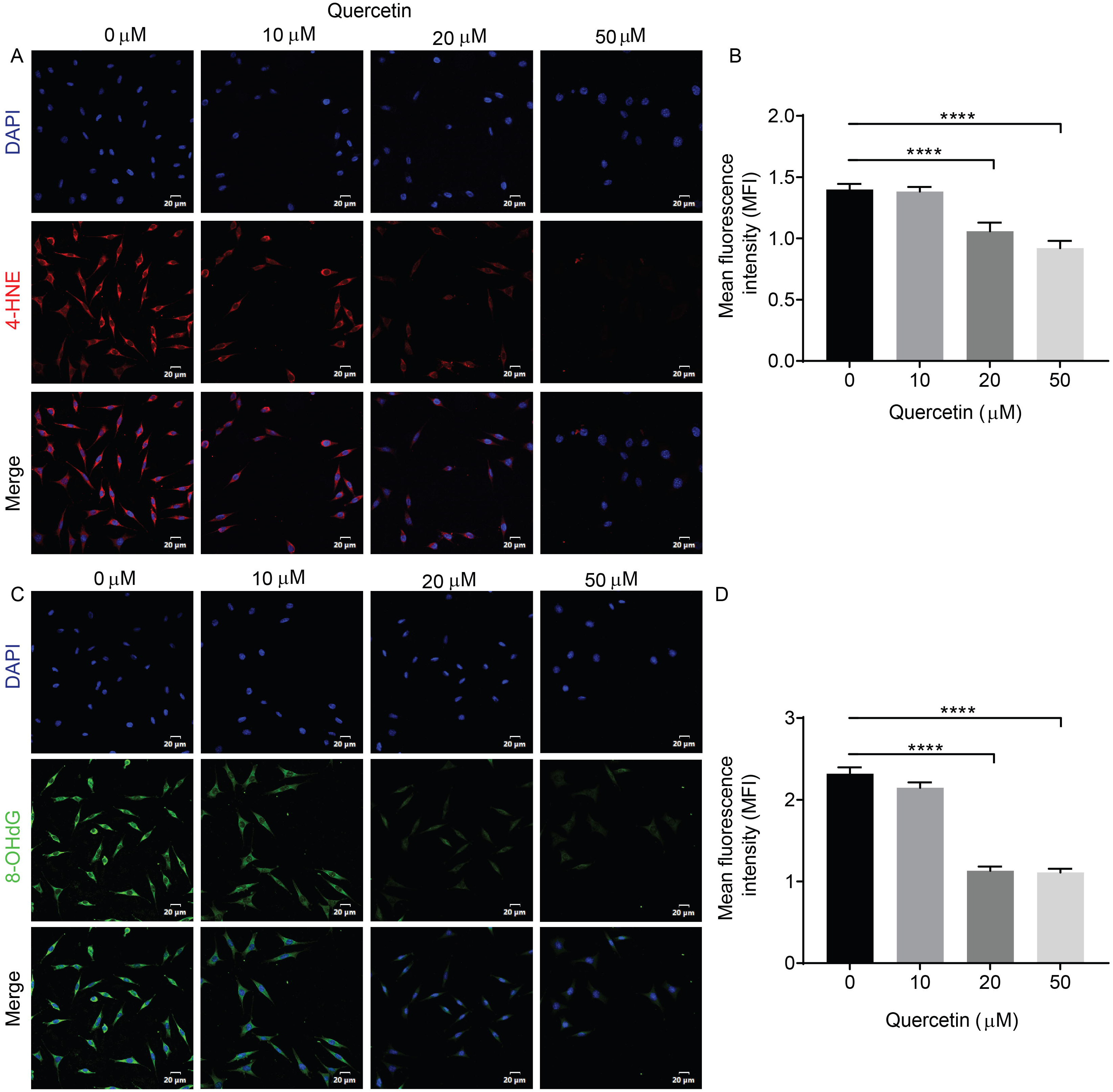
Quercetin attenuates the increase in the level of lipid peroxidation and prevents DNA oxidation in N2a/APP cells. A, Representative confocal images of the immunoreactivity of 4-hydroxynonenal (4-HNE, red) in N2a/APP cells without or with quercetin treatment at the concentration of 10 μM, 20 μM, 50 μM. Nuclei are stained blue with DAPI; scale bar = 20 μm; B, Quantitation of the fluorescence intensity (n = 3). A total of 100 cells from each group were analysed. **** p < 0.0001, one-way ANOVA, Dunnett’s multiple comparison. C, Representative confocal images of anti-8-hydroxy-2’-deoxyguanosine (8-OHdG) immunofluorescent staining (green) in N2a/APP cells without or with quercetin treatment at the concentration of 10 μM, 20 μM, 50 μM. Nuclei are stained blue with DAPI. Scale bar = 20 μm; D, Quantification of the mean fluorescence intensity (n = 3). A total of 100 cells from each group were analysed. **** p < 0.0001, one-way ANOVA, Dunnett’s multiple comparison.

## 4. Discussion

Here, we observed that quercetin downregulated the ERK1/2 pathway and upregulated the AKT pathways, while it protected N2a cells against the neurotoxicity of abnormal APP by reducing the generation of ROS, lipid peroxidation and DNA oxidation, as well as re-establishing mitochondrial function.

In our current study, we examined the effects of quercetin on murine neuroblastoma N2a cells stably expressing human Swedish mutant APP, a well-characterized cellular model of AD. We found higher levels of APP and the amyloidogenic pathway, consistent with previous reports (Butterfield and Boyd-Kimball, 2018; Cheignon et al., 2018). We observed that quercetin reduced cellular APP expression in N2a/APP cells. Our data are supported by previous *in vivo* studies in which quercetin treatment induced a significant decrease in the levels of C-terminal APP fragments and in the Aβ40 and Aβ42 levels in the hippocampus of the quercetin-treated 3×Tg-AD mice (Sabogal-Guaqueta et al., 2015).

We found that quercetin significantly decreased the expression of phosphorylated ERK1/2 in N2a/APP cells, which is consistent with a previous report describing quercetin protecting against Aβ-induced toxicity in SH-SY5Y cells by inhibiting the ERK1/2 signaling pathway (Ansari et al., 2009; Shi et al., 2009). We found that quercetin increased the expression of phosphorylated-AKT in N2a/APP cells. Previous study by Tangsaengvit et al. has reported that lower concentration of quercetin (1 nM) significantly increased the amount of neuritis and neurite length via the activation of P13K/AKT pathway (Tangsaengvit et al., 2013). However, mechanistically, quercetin activates or inactivates the expression of several additional signaling cascades, i.e., mitogen-activated protein kinases (MAPKs), mitophagy activation, sirtuin 1 (SIRT1) and nuclear factor erythroid 2-related factor 2/heme oxygenase-1(Nrf2/HO-1) pathway (Cen et al., 2022; Cui et al., 2022; Spencer et al., 2003; Xie et al., 2022). Further studies are needed to elucidate the neuroprotective effect of quercetin in vivo with a more systematic approach.

Our evidence demonstrated that N2a/APP cells generate more intracellular ROS, lipid peroxidation (4-HNE) and oxidation of DNA (8-OHdG). We found that pretreatment of N2a/APP cells with quercetin suppressed ROS generation, reduced the formation of a lipid peroxidation product (4-HNE), decreased the level of a DNA damage marker (8-OHdG) and attenuated mitochondrial dysfunction. Oxidative stress is caused by the accumulation of ROS in neurons and contributes to the development of neurodegeneration (Niedzielska et al., 2016; Su et al., 2008). Accumulated oxidative stress can cause cellular damage, DNA repair system impairment and mitochondrial dysfunction leading to oxidative imbalance, and thus plays an important role in AD progression (Islam, 2017; Mecocci et al., 2018; Su et al., 2008). Abberant Aβ accumulation to induce mitochondrial impairment and leads to oxidative stress and power failure (Butterfield and Boyd-Kimball, 2018; Cheignon et al., 2018). The oxidative imbalance caused by Aβ increases ROS production and results in a dissipation of ΔΨm, indicating mitochondrial dysfunction, as well as the levels of lipid peroxidation and oxidation of DNA (Butterfield and Boyd-Kimball, 2018; Richter, 1995). Ansari et al. found that quercetin protected against Aβ_1-42_-induced cell toxicity in primary hippocampal neurons and that pretreatment with quercetin drastically lowered Aβ_1-42_-induced neurotoxicity, lipid peroxidation, protein oxidation and apoptosis (Ansari et al., 2009). Most previously reported interventions designed potentially to protect against Aβ rely on antioxidants to relieve ROS-induced oxidative stress.

Quercetin have demonstrated the effect on other cellular types in addition to neurons; such as alleviateing Aβ-associated oligodendrocyte progenitor cell senescence and cognitive deficits in APP/PS1 model of AD amyloidosis (Zhang et al., 2019), involved in microglia associated reduction of the accumulation of Aβ plaques and inflammation in AD model mice (Sabogal-Guaqueta et al., 2015; Wang et al., 2022). Further studies are needed to employ a co-culture model with neurons, astrocytes and microglia to study the effect of quercetin.

## 5. Conclusion

In conclusion, we demonstrate that quercetin inhibits abnormal APP expression on N2a/APP cells by regulating mitochondrial dysfunction and reducing the generation of ROS, lipid peroxidation, and DNA oxidation. Furthermore, quercetin may exert these effects through the inhibition of the ERK1/2 pathways, as well as activation of the AKT signaling pathway. These results may offer new insight to better understand the defensive mechanism of quercetin against abnormal APP neurotoxicity.

## Supporting information

westernblot for fig 2E-H

westernblot for fig 2A-D

westernblot for fig 1

## Author Contributions

ZT, RN, XLQ contributed to the conception and study design. MG, YQP and TZ performed the experiments. ZT, MG and YX contributed to data collection and data analysis. RN and ZT interpreted the data. XLQ, GM, RN, and ZT wrote the manuscript. All authors approved the manuscript before submission.

