## Supplementary table 1 for "Quercetin reduces APP expression, oxidative stress and mitochondrial dysfunction in the N2a/APPswe cells via ERK1/2 and AKT pathways"

**Supplementary Table 1 List of chemicals, antibodies and kits used in the study**

| **Product** | **Catalogue no** | **Company** |
| --- | --- | --- |
| Quercetin | #117-39-5 | MCE, USA |
| Dimethyl sulfoxide | #D2650 | Sigma, USA |
| Proteasehemmer-Cocktail | #P8340 | Sigma, USA |
| Antibody against total-ERK | #4695 | Cell Signaling Technology, USA |
| Antibody against phospho-ERK | #4370 | Cell Signaling Technology, USA |
| Antibody against total-ATK | #9272 | Cell Signaling Technology, USA |
| Antibody against phospho-AKT | #4060 | Cell Signaling Technology, USA |
| Antibody against APP | #ab32136 | Abcam, USA |
| 4-Hydroxynonenal | #ab46545 | Abcam, USA |
| 8-Hydroxyguanosine | #ab62623 | Abcam, USA |
| Antibody against GAPDH | #GTX100118 | Genetex, USA |
| DCFH-DA | #S0033S | Beyotime, China |
| JC-1 | #C2006 | Beyotime, China |
| PVDF membranes | #ISEQ00010 | Millipore, USA |
| Bradford kit | #5000205 | Bio-Rad, USA |
| Dulbecco’s modified Eagle’s medium | #11965092 | Gibco, USA |
| Fetal bovine serum | #10100147 | Gibco, USA |
| Penicillin‒streptomycin | #SV30010 | HyClone, USA |
| Phosphate-buffered saline | #02-020-1A | Biological Industries, Israel |
| RIPA cell lysis buffer | #R0010 | Solarbio, China |
| Cell counting kit-8 | #HY-K0301 | MedChemExpress LLC, China |
| DAPI Fluoromount-G | #0100-20 | SouthernBiotech, USA |
| SDS polyacrylamide gels | #abs9382 | Absin, China |
| Non-fat milk | #CN7861 | Coolaber, China |
| Tris-buffered saline | #G0001 | Servicebio, China |
| Tween-20 | #T8220 | Solarbio, China |
| Triton X-100 | #T8200 | Solarbio, China |
| Goat anti-Mouse IgG (H+L), HRP | #31430 | Thermo Fisher Scientific, USA |
| Goat anti-Rabbit IgG (H+L), HRP | #31460 | Thermo Fisher Scientific, USA |
| Alexa Fluor 488 | #A21202 | Thermo Fisher Scientific, USA |
| Alexa Fluor 546 | #A10040 | Thermo Fisher Scientific, USA |

APP, amyloid precursor protein; DAPI, 4′,6-diamidino-2-phenylindole; DCFH-DA, 2’,7’-dichlorofluorescein diacetate; ERK, extracellular signal-regulated kinase; GAPDH, glyceraldehyde 3-phosphate dehydrogenase; HRP, horseradish peroxidase; PMSF, phenylmethylsulfonyl fluoride; SDS, Sodium dodecyl sulfate; JC-1, 5,5,6,6’-tetrachloro-1,1’,3,3’-tetraethylbenzimi-dazoylcarbocyanine iodide; 4-HNE, 4-hydroxynonenal; 8-OHdG, 8-hydroxyguanosine;
