## Supplementary figures and images for "Quercetin reduces APP expression, oxidative stress and mitochondrial dysfunction in the N2a/APPswe cells via ERK1/2 and AKT pathways"

### westernblot for fig 1

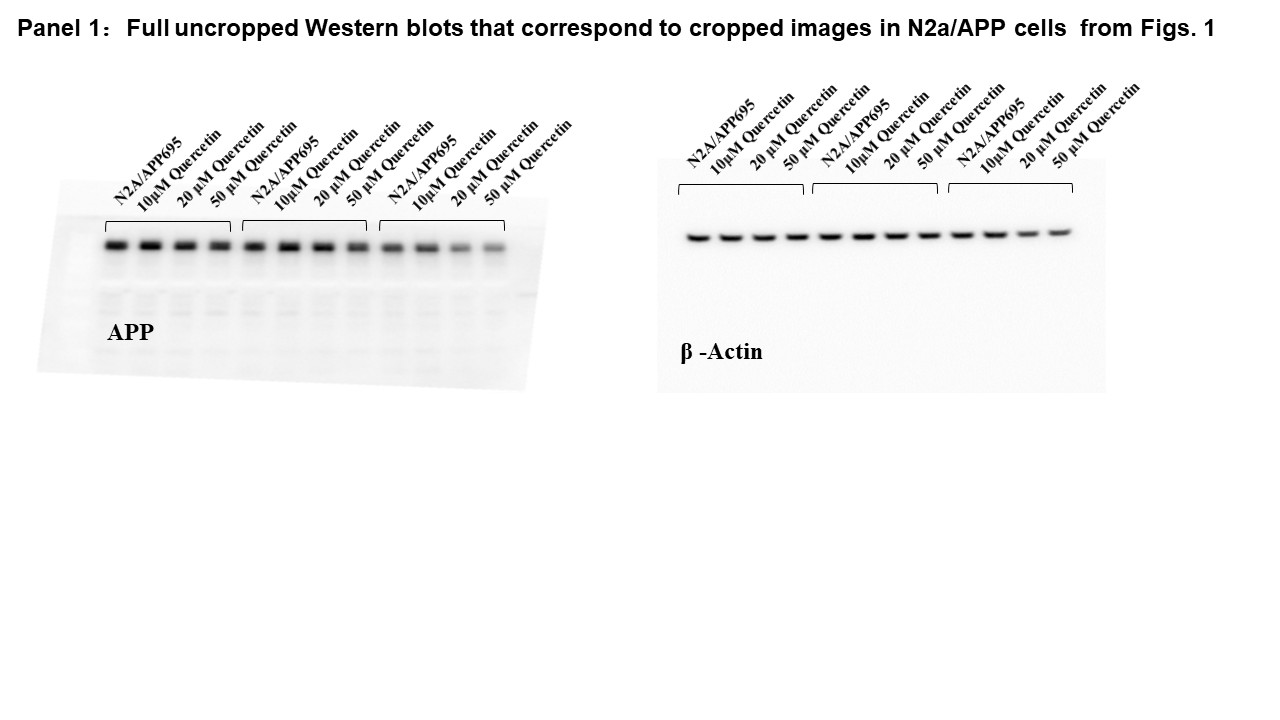

### westernblot for fig 2A-D

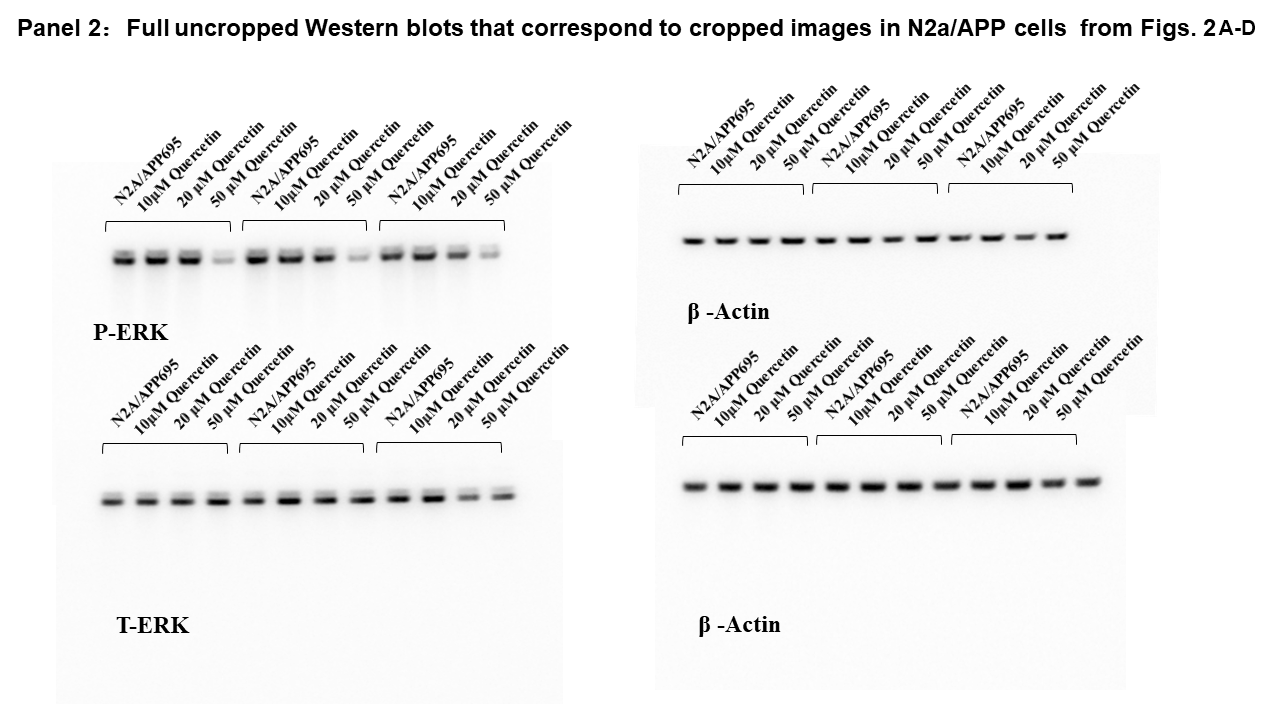

### westernblot for fig 2E-H

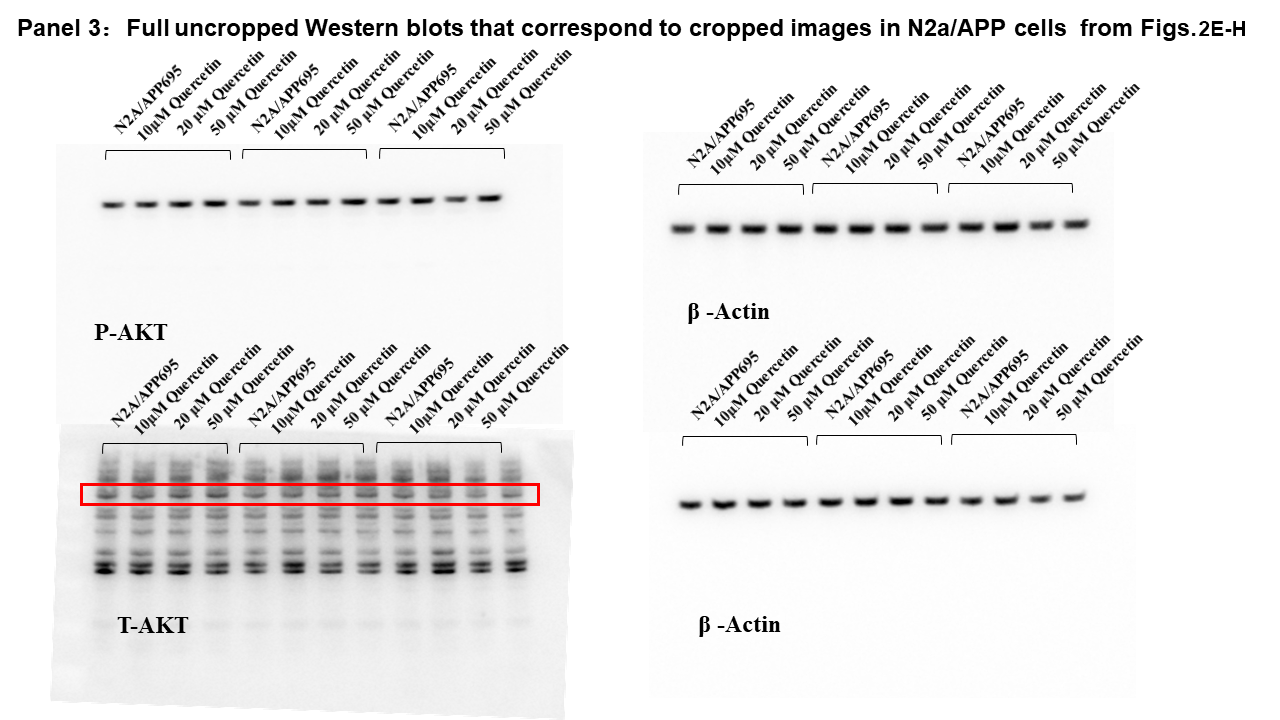
